# ITK signaling regulates a switch between T helper 17 and T regulatory cell lineages via a calcium-mediated pathway

**DOI:** 10.1101/2023.04.01.535229

**Authors:** Orchi Anannya, Weishan Huang, Avery August

**Author notes:** Address correspondence to: Avery August Department of Microbiology & Immunology College of Veterinary Medicine 930 Campus Rd Cornell University Ithaca, NY14853.

## Abstract

The balance of pro-inflammatory T helper type 17 (Th17) and anti-inflammatory T regulatory (Treg) cells is crucial in maintaining immune homeostasis in health and disease conditions. Differentiation of naïve CD4^+^ T cells into Th17/Treg cells is dependent upon T cell receptor (TCR) activation and cytokine signaling, which includes the kinase ITK. Signals from ITK can regulate the differentiation of Th17 and Treg cell fate choice, however, the mechanism remains to be fully understood. We report here that in the absence of ITK activity, instead of developing into Th17 cells under Th17 conditions, naïve CD4^+^ T cells switch to cells expressing the Treg marker Foxp3 (Forkhead box P3). These switched Foxp3^+^ Treg like cells retain suppressive function and resemble differentiated induced Tregs in their transcriptomic profile, although their chromatin accessibility profiles are intermediate between Th17 and induced Tregs cells. Generation of the switched Foxp3^+^ Treg like cells was associated with reduced expression of molecules involved in mitochondrial oxidative phosphorylation and glycolysis, with reduced activation of the mTOR signaling pathway, and reduced expression of BATF. This ITK dependent switch between Th17 and Treg cells was reversed by increasing intracellular calcium. These findings suggest potential strategies for fine tune the TCR signal strength via ITK to regulate the balance of Th17/Treg cells.

## INTRODUCTION

Immune function mediated by T helper (Th) cells is critical for the development of effective barriers to pathogens and environmental toxins. Distinct functions are mediated by specific lineages of T helper cells that in general have pro-inflammatory function such as that mediated by Th1, Th2, Th9 and Th17 lineages, and anti-inflammatory function such as that mediated by the T regulatory (Treg) and T regulatory type 1 (Tr1) lineages. These distinct T helper cells lineages are characterized by expression of lineage specific transcription factors and the effector cytokines they produce that mediate immune response (*1*). Th17 cells are characterized by expression of the transcription factor Rorγt (RAR-related orphan receptor gamma), and produce effector cytokines IL17A, IL17F, IL21, IL22 in response to infection with extracellular bacteria, parasites and allergens (*1*). Tregs are characterized by expression of the transcription factor Foxp3 (Forkhead box P3) and produce effector cytokines such as IL10 and TGFβ, among other effectors that can suppress function of Th1, Th2, Th9 and Th17 cells among other immune cells (*1*).

Commitment of naïve CD4^+^ T cells to generally pro-inflammatory Th17 effector fates, or to generally anti-inflammatory Foxp3^+^ Treg fates, have been shown to be dependent on the strength of TCR signaling and the cytokines present in the microenvironment (*2-6*). Key events include triggering of TCR upon interaction with antigen/MHC complexes and subsequent recruitment of the Src family of kinases Lck/Fyn, and Syk family of kinase ZAP-70 (zeta chain associated kinase), which phosphorylates adaptor proteins LAT (linker for activation of T cell) and SLP76 (SH3 containing lymphocyte protein) (*7, 8*). This is followed by assembly of a proximal signaling complex that includes the recruitment of ITK (IL-2 inducible T cell kinase) and SLP76 (*7, 8*). ITK functions in part by activating PLCγ (phospholipase Cγ) to generate effector molecules IP3 (inositol triphosphate) and DAG (diacylglycerol) (*7-9*). This allows activation of MAPK (mitogen activated protein kinase) cascades, enhanced cytosolic calcium concentrations required to activate NFAT (nuclear factor of activated T cells) and initiation of Akt dependent mTOR (mammalian target of rapamycin) (*8*). TCR signal strength has been shown to be regulated by ITK (*8*). While the initial stages of T cell activation is dependent on TCR activation and presence of select cytokines in the microenvironment, once activated the T cells undergo further metabolic changes unique to each T cell subset (*10*). These metabolic changes allow maintenance and function as select T cell subsets, for example Th17 cells are highly dependent on mitochondrial oxidative phosphorylation and glycolysis (*11, 12*). These metabolic changes in turn regulate expression of key molecules that allow function of these T cell subsets as pro-inflammatory Th17 or anti-inflammatory Treg cells (*10*).

Given the critical role of ITK in regulating development and function of T cell lineages, several studies have investigated the function of ITK in the development of difference immune responses. We and others have also shown that in naïve CD4^+^ T cells, the absence of ITK, or ITK activity, impairs differentiation into Th2, Th17 and Tr1 lineages while enhancing differentiation into Th1 and Treg lineages (*13-18*). In particular, TCR signal strength traveling via ITK has been shown to play an important role determining Th17 versus Treg lineage commitment (*14, 17, 19*). However, whether inhibition of ITK results a similar regulation and the mechanisms by which this dichotomy is controlled is unclear.

Here, we investigate this by utilizing ITK inhibitors and allele sensitive ITK (ITK*as*) expressing mice allowing selective inhibition by ITK*as* selective inhibitor compounds (*16, 20-23*). We show that under conditions that promote Th17 differentiation, inhibition of ITK suppresses naïve CD4^+^ T cells differentiation to Th17 lineage, and instead leads to the generation of Foxp3^+^ Treg-like cells. The resultant population of Foxp3^+^ Treg-like cells express markers associated with Tregs, effectively retain expression of Treg transcription factor Foxp3, and demonstrate effective suppression of responding T cell proliferation. Furthermore, the switched Foxp3^+^ Treg-like cells have Treg-like transcriptomic and chromatin accessibility profile. The resultant Foxp3^+^ Tregs resemble iTregs by displaying reduced expression of key markers involved in mitochondrial oxidative phosphorylation and the glycolytic pathways. This overall reduction in oxidative phosphorylation was associated with reduced phosphorylation of mTOR and downstream substrate S6K (ribosomal S6 kinase), which may act to prevent expression of the Th17 pioneer transcription factor BATF (basic leucine zipper ATF-like transcription factor). Finally, we show that bypassing the TCR signal to increase calcium signaling prevents this ITK dependent switch response. Our results indicate that ITK signals can tune the balance between Th17 cells and Tregs.

## RESULTS

### ITK controls a switch between Th17 differentiation and Foxp3^+^ Treg-like cells when naïve CD4^+^ T cells are activated under Th17 differentiation conditions

In the absence of ITK activity, Th17 differentiation is inhibited, and in *Itk^-/-^* T cells, the absence of ITK results in reduced Th17 differentiation, and the appearance of Foxp3^+^ Treg cells (*14-17*). However, whether this switch from Th17 to Treg differentiation occurs when ITK kinase activity is inhibited is unclear. To determine the role of ITK activity in the development of Foxp3^+^ T regulatory cells under Th17 conditions, we stimulated naïve WT, *Itk^-/-^* or ITK*as* T cells isolated from IL17A-GFP/Foxp3-RFP reporter mice, under Th17 culture conditions. We found that the absence of ITK, or inhibition of ITK*as* results in the inhibition of Th17 cell differentiation, along with the appearance of Foxp3^+^ Treg-like cells (**Fig. 1a**). Similarly, stimulation of naïve WT IL17A-GFP/Foxp3-RFP reporter cells under Th17 culture conditions in the presence of the covalent small molecule ITK inhibitor (*23*) (CPI-818) resulted in dose dependent inhibition of Th17 cell differentiation, along with the appearance of Foxp3^+^ Treg-like cells, with similar results observed when naïve ITK*as* T IL17A-GFP/Foxp3-RFP reporter cells were used along with MBPP1 which we have previously shown selectively inhibits the ITK*as* isoform (*16*) (**Fig. 1b,c**). This effect was due to specific inhibition of ITK and not potential off-target effects since the covalent small molecule ITK inhibitor (CPI-818) does not affect Th17 differentiation of naïve ITK*as* T cells, nor does MBPP1 affect Th17 differentiation of naïve WT T cells (**Supplemental Fig. 1**). We refer to these Foxp3^+^ Treg-like cells generated under conditions of ITK inhibition as *Switched Foxp3^+^ Treg-like* cells.

**Fig. 1.**
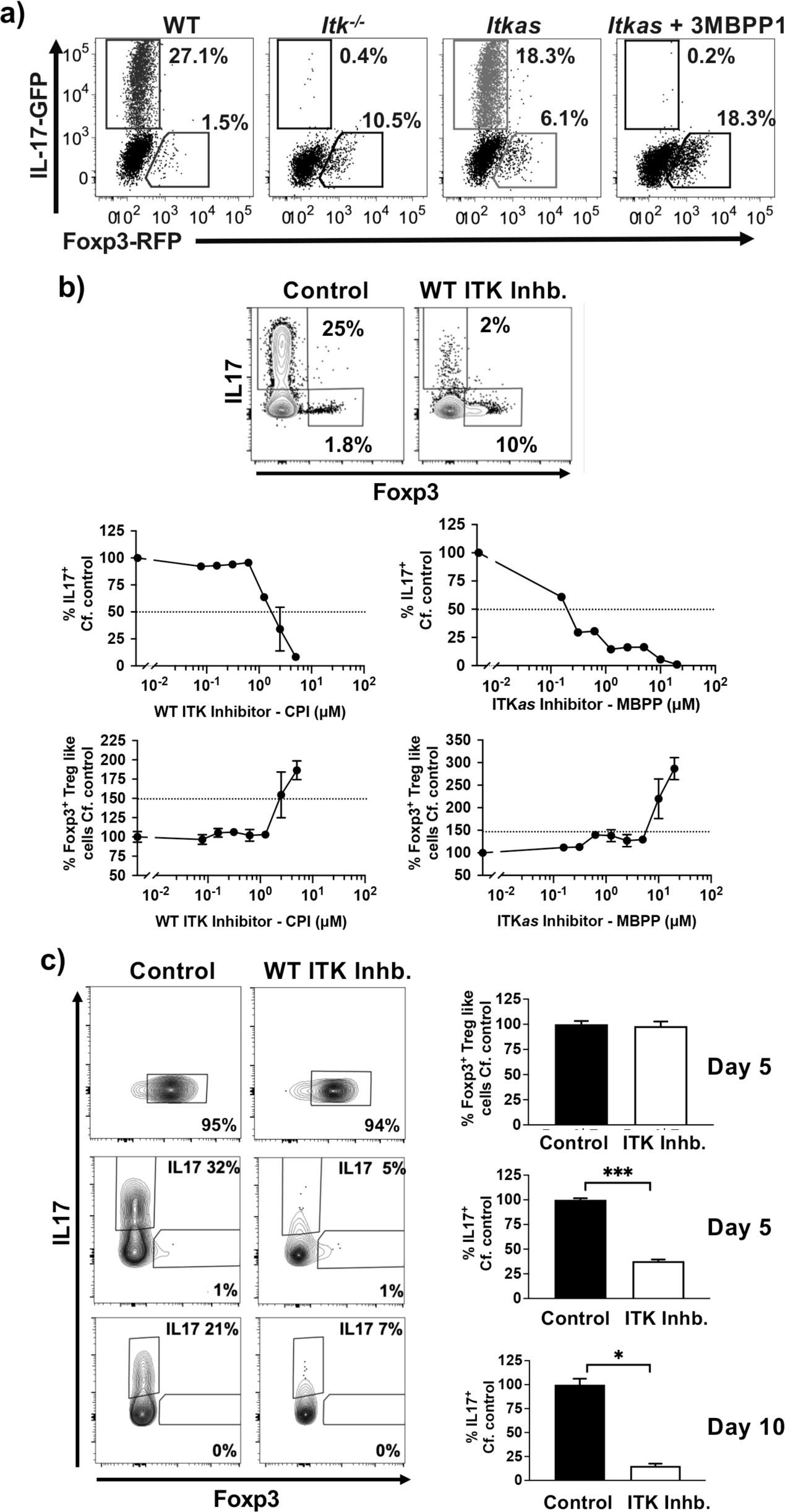
ITK controls a switch between Th17 differentiation and Foxp3^+^ Treg-like cells when naïve CD4^+^ T cells are activated under Th17 differentiation conditions. (**a**) Naïve WT, *Itk^-/-^* or ITK*as* IL17A-GFP/Foxp3-RFP CD4^+^ T cells were activated under Th17 differentiation conditions (anti-CD3/28, IL6 and TGFβ) in presence of ITK*as* inhibitor 2MBPP1 or DMSO control as indicated, followed by flow cytometric analysis for percentage of GFP^+^/IL17^+^ cells and RFP^+^/Foxp3^+^ Treg-like cells. (**b**) Top: Naïve WT IL17A-GFP/Foxp3-RFP CD4^+^ T cells were activated under Th17 differentiation conditions in presence of ITK inhibitor CPI-818 or DMSO control, followed by quantification of percentage of GFP^+^/IL17^+^ cells and RFP^+^/Foxp3^+^ Treg-like cells. Bottom: Naïve WT (left) or ITK*as* (right) IL17A-GFP/Foxp3-RFP CD4^+^ T cells were activated under Th17 differentiation conditions in presence of ITK inhibitor CPI-818 (left) ITK*as* inhibitor 3MBPP1 (right) or DMSO control, followed by quantification of percentage of GFP^+^/IL17^+^ cells and RFP^+^/Foxp3^+^ Treg-like cell. Mean ± SEM, one-way ANOVA was performed for statistical significance where * p ≤ 0.05, ** p ≤ 0.005, *** p ≤ 0.001 and **** p ≤ 0.0001, 3 independent experiments. (**c**) Switched Foxp3^+^ Treg-like cells generated under Th17 differentiation conditions in presence of ITK inhibitor, or *in vitro* generated Th17 cells, were then further reactivated under Th17 differentiation conditions in presence of ITK inhibitor CPI-818 or DMSO control for a duration of 5 days (Switched Foxp3^+^ Treg-like cells and Th17 cells) or 10 days (Th17 cells). Mean ± SEM, Student’s T test was performed for statistical significance where * p ≤ 0.05, ** p ≤ 0.005, 3 independent experiments.

### ITK inhibition does not switch already differentiated Th17 cells to Foxp3^+^ Treg-like cells

To determine whether inhibition of ITK can induce already differentiated Th17 cells to become switched Foxp3^+^ Treg-like cells, we generated *in vitro* both the switched Foxp3^+^ Treg-like cells and Th17 cells, sorted them (by GFP+/RFP-expression), and further cultured the sorted cells under Th17 differentiation conditions in presence of the covalent ITK inhibitor. While we there was no change in the switched Foxp3^+^ Treg-like cells, inhibiting ITK led to a reduction in differentiated Th17 cells suggesting that ITK activity is required for the maintenance of these cells. However, inhibiting ITK in these Th17 cells did not lead to the development of Foxp3^+^ Treg-like cells under these conditions (**Fig. 1c**). This data suggests that under Th17 differentiation conditions, inhibition of ITK switches Th17 differentiation to Foxp3^+^ Treg-like cells, but is not able to do so once the cells have already differentiated to the Th17 fate.

### Early events following TCR activation determines the ability of ITK to tune a switch from Th17 to Foxp3^+^ Treg-like cells

To determine if early or late molecular events influence this switch to Foxp3^+^ Treg-like cells upon ITK inhibition, naïve CD4^+^ T cells were activated under Th17 differentiation conditions, followed by addition of the ITK inhibitor after 1, 2, 3, or 4 days post initiation of the culture. We find that inhibition of ITK time dependently (after up to 2 days) blocks Th17 differentiation (**Fig. 2a**). Analysis of the generation of switched Foxp3^+^ Treg-like cells also showed a time dependent relationship between ITK inhibition, and the generation of these cells, although optimal generation required early ITK inhibition, as ITK inhibition at later time points beyond day 1 is less effective in inducing the switch, with some Foxp3^+^ Treg-like cell generation upon ITK inhibition on day 2, and no significant effect upon ITK inhibition between days 3 to 5 (**Fig. 2a**).

**Fig. 2.**
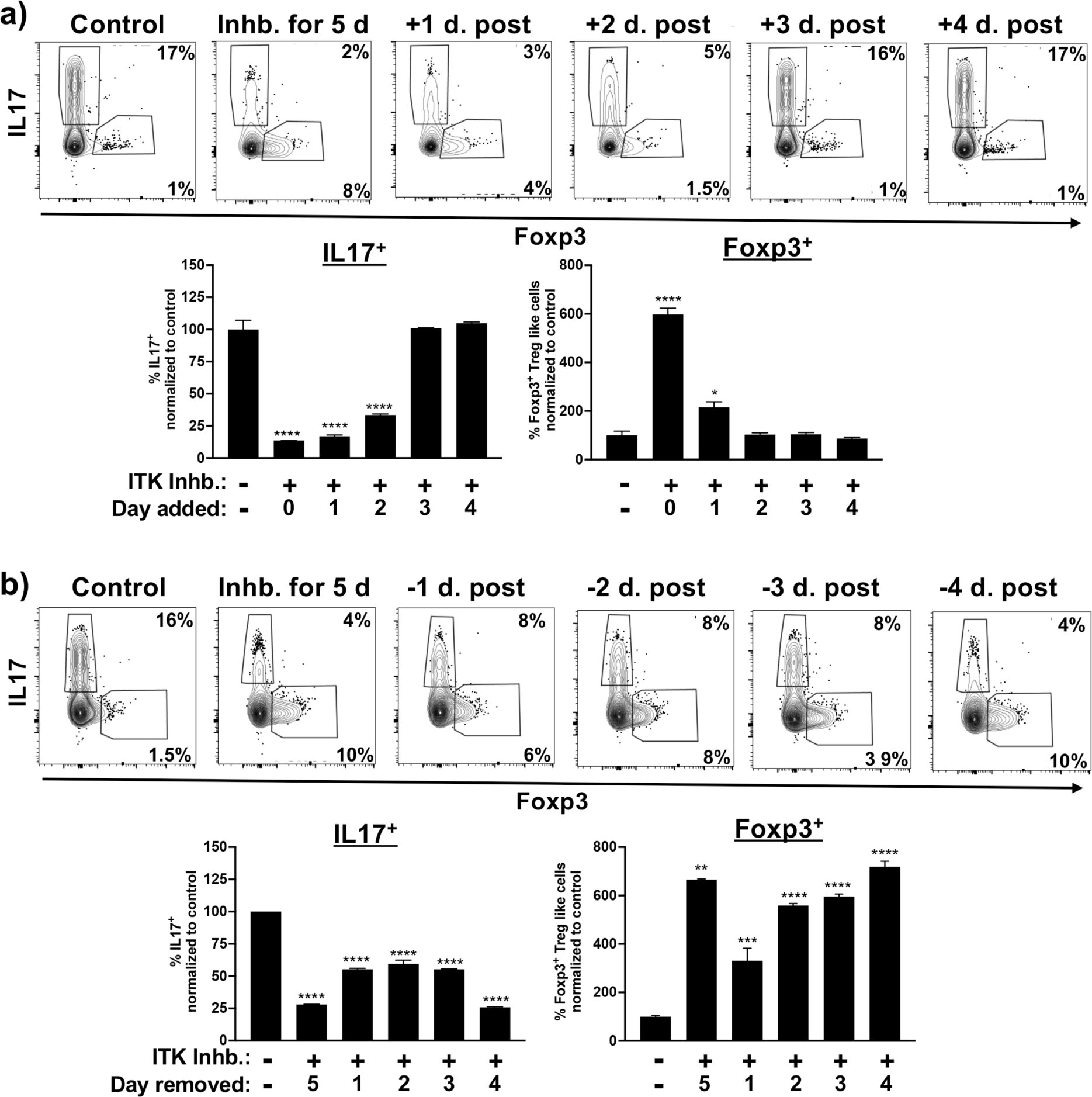
Early ITK signals are required for ITK mediated switch between Th17 and switched Foxp3^+^ Treg-like cells under Th17 differentiation conditions. (**a**) Naïve WT IL17A-GFP/Foxp3-RFP CD4^+^ T cells were activated under Th17 differentiation conditions followed by addition of ITK inhibitor CPI-818 after 1, 2, 3 or 4 days of culture, followed by analysis on day 5. (**b**) Naïve WT IL17A-GFP/Foxp3-RFP CD4^+^ T cells were activated under Th17 differentiation conditions in the presence of the ITK inhibitor CPI-818, followed by removal of inhibitor after 1, 2, 3 or 4 days of culture, with analysis on day 5. Mean ± SEM, one-way ANOVA was performed for statistical significance where * p ≤ 0.05, ** p ≤ 0.005, *** p ≤ 0.001 and **** p ≤ 0.0001, 2 independent experiments.

Next, we wanted to determine the time frame of ITK inhibition that results in this switch response. Using a similar approach, naïve CD4^+^ T cells were activated under Th17 differentiation conditions, followed by removal of the ITK inhibitor after 1, 2, 3, or 4 days. Removal of ITK inhibitor after 1 day led to an ∼50% inhibition of Th17 cell differentiation, although, with the greatest effect observed with 5 days of ITK inhibition (**Fig. 2b**). Analogously, the generation of switched Foxp3^+^ Treg-like cells was observed when ITK was inhibited for as little as 1 day (∼40% of maximal generation), and close to maximum after 3 days of ITK inhibition (**Fig. 2b**). Taken together this suggests that early events following TCR activation determines the switch from Th17 to switched Foxp3^+^ Treg-like cells upon ITK inhibition.

### Switched Foxp3^+^ Treg-like cells resemble true Foxp3^+^ Tregs and can suppress naïve T cell proliferation *in vitro*

In addition to expression of Foxp3, Treg cells express a number of cell surface molecules including CD25, CTLA4 and PD1 (*1*). To determine if the switched Foxp3^+^ Treg-like cells generated upon ITK inhibition of naïve CD4^+^ T cells cultured under Th17 conditions, resemble true Tregs, we compared the expression of these cell surface markers by flow cytometry. We find that all Treg subsets (*in vitro* generated induced or iTregs, thymic derived natural or nTregs and extra-thymic peripheral or pTregs) as well as the switched Foxp3^+^ Treg-like cells express CD25, CTLA4 and PD1, with switched Foxp3^+^ Treg-like cells closely resembling the iTregs in their level of expression of these markers (**Fig. 3a**). Subsequently we investigated if the switched Foxp3^+^ Treg-like cells are able to suppress proliferation of CFSE-labeled naïve CD4^+^ T cells when cocultured *in vitro*. In the absence of coculture with Tregs, responding naïve T cells are able to undergo multiple rounds of cell division when activated by anti-TCR antibodies (as determined by CFSE dye dilution). However, when cocultured in presence of iTregs, this proliferation is suppressed (**Fig. 3b**). In presence of the switched Foxp3^+^ Treg-like cells, similar to the iTregs, this proliferation of responding T cells is also suppressed (**Fig. 3b**). Therefore, the switched Foxp3^+^ Treg-like cells, express surface markers and display suppressive function similar to the iTregs.

**Fig. 3.**
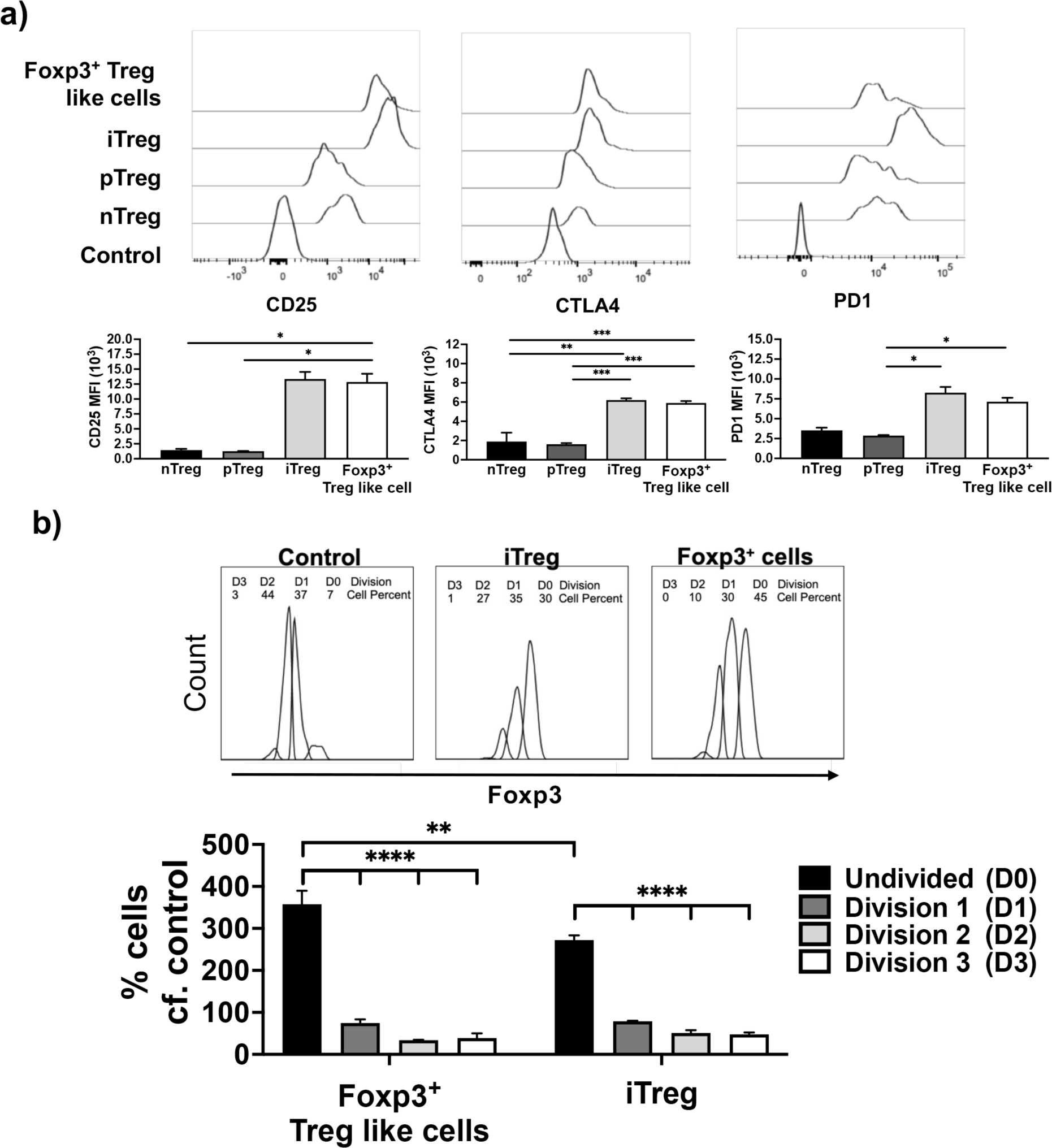
Switched Foxp3^+^ Treg-like cells generated under conditions of ITK inhibition have a cell surface phenotype and suppressive function similar to iTregs. (**a**) Foxp3^+^ Treg-like cells that are generated from WT IL17A-GFP/Foxp3-RFP CD4^+^ T cells activated under Th17 differentiation conditions in presence of ITK inhibitor CPI-818, and compared to iTregs, pTregs and nTregs for expression of select Treg markers. Expression represented as mean fluorescence intensity. (**b**) CD45.2 switched Foxp3^+^ Treg-like cells (generated in absence of ITK activity under Th17 conditions) and iTregs were sort purified, followed by co-culture with CFSE labelled CD45.1 naïve CD4^+^ T cell responders. Representative CFSE plots (top). Percentage suppression of proliferation of naïve CD4^+^ T cell responders by Foxp3^+^ Treg-like cells and iTregs was quantified (bottom). Mean ± SEM for percentage of cells undergoing division. One-way ANOVA performed for statistical significance where * p ≤ 0.05, ** p ≤ 0.005, *** p ≤ 0.001 and **** p ≤ 0.0001, 3 independent experiments.

### Switched Foxp3^+^ Treg-like cells have a transcriptome that resemble iTregs

We next investigated the transcriptomic profile of the switched Foxp3^+^ Treg-like cells, comparing then to *in vitro* derived Th17 cells and iTregs, using RNA-Seq analysis. Principal component analysis and heatmap analysis displaying expression of all transcripts, indicated that the switched Foxp3^+^ Treg-like cells closely resemble iTregs compared to Th17 cells (**Fig. 4a, b**). Using volcano plots to identify genes differentially expressed, we find that Th17 cytokine Il17 and the critical Th17 transcription factor Rorc were downregulated in switched Foxp3^+^ Treg-like cells compared to Th17 cells (**Fig. 4c**). Furthermore, Foxp3, and Treg-related genes such as tgfbr1 and tgfbr2, as well as other transcription factors smad3 and Ikzf2 were found to be upregulated in switched Foxp3^+^ Treg-like cells compared to Th17 cells (**Fig. 4c**). Notably, the switched Foxp3^+^ Treg-like cells expressed lower levels of Treg related genes foxp3, nrp1, tgfb1 and tgfbr1, although higher levels of tgfb3 (**Fig. 4c**).

**Fig. 4.**
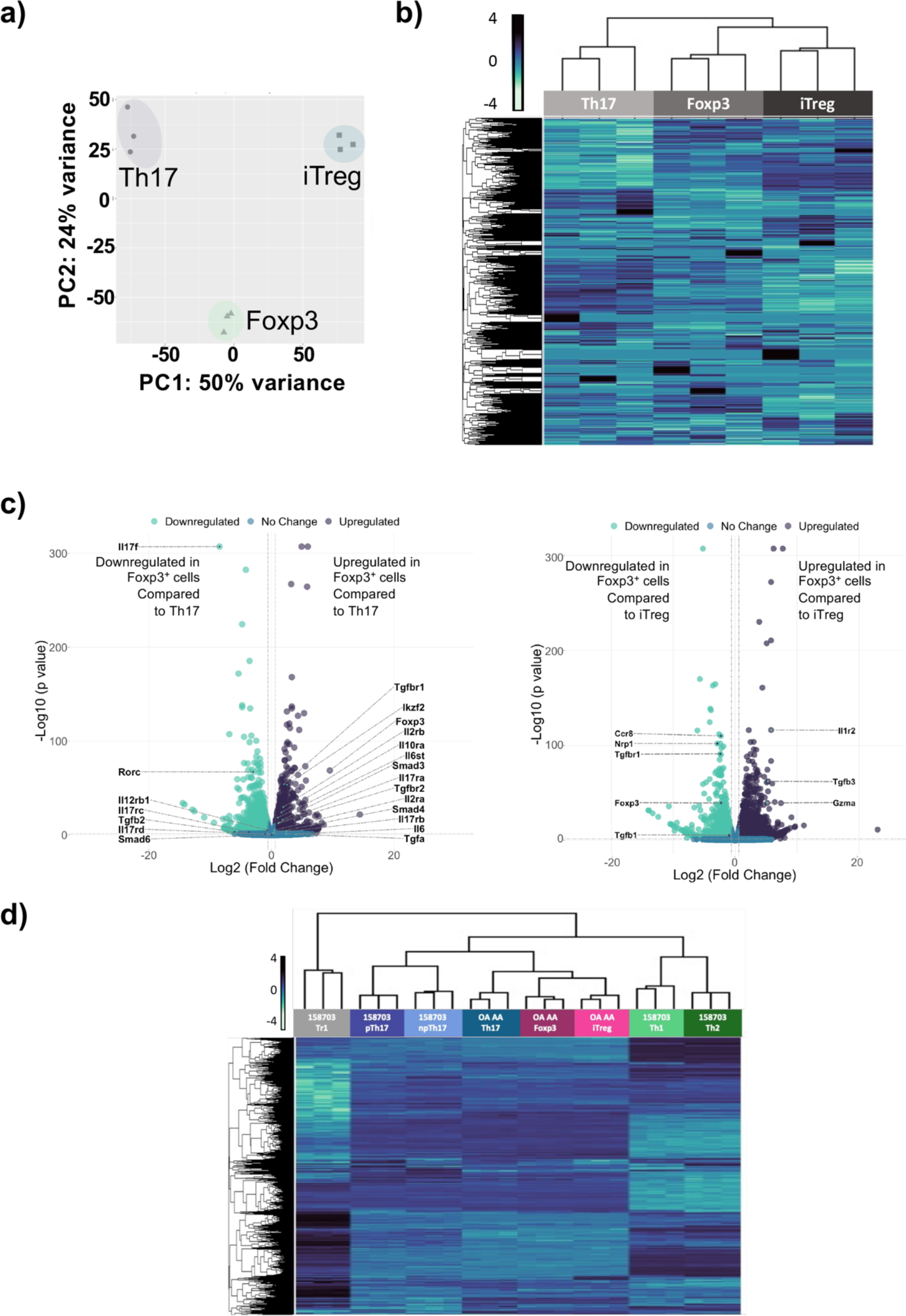
Switched Foxp3^+^ Treg-like cells generated under ITK inhibition have a transcriptomic profile similar to iTregs. Switched Foxp3^+^ Treg-like cells were compared to *in vitro* generated Th17 and iTregs by RNA-Seq analysis. The transcriptome was compared via (**a**) PCA analysis, (**b**) heatmap of global gene expression, or **(c**) volcano plot of differentially expressed genes. N-3 for each subset. (**d**) Switched Foxp3^+^ Treg-like cells were compared with GEOdata set from GSE158703 by heatmap.

To compare the transcriptomes of switched Foxp3^+^ Treg-like cells with other T cell subsets, we compared the transcriptomes to transcriptomic data of Th1, Th2, pathogenic pTh17 cells generated in presence of IL1, IL6 and IL23, non-pathogenic npTh17 cells generated in presence of IL6 and TGFβ and Tr1 cells (GSE158703)(*24*). We find that the transcriptome of switched Foxp3^+^ Treg-like cells, Th17 and iTreg subsets we generated cluster closer to the npTh17 and pTh17 subsets, and further from Th1 and Th2 subsets, with Tr1 cells being furthest in the cluster comparison (**Fig. 4d**).

### Switched Foxp3^+^ Treg-like cells generated under ITK inhibition have a chromatin profile distinct from both Th17 and iTreg cell subsets

We also analyzed the accessibility of the chromatin in switched Foxp3^+^ Tregs by ATAC-Seq, comparing them to Th17 and iTreg cell subsets. Analysis by PCA and heatmaps revealed that similar to the transcriptome, switched Foxp3^+^ Treg-like cells have chromatin profiles that resemble iTreg compared to Th17 cells (**Fig. 5a,b**). Using volcano plots to compare differentially open and closed chromatin regions, we find that the chromatin region of genes associated with Th17 cells (e.g., Rora, Stat3, Il17, Foxp1) displayed reduced accessibility in switched Foxp3^+^ Treg-like cells compared to Th17 cells (**Fig. 5c**). By contrast, the chromatin regions of genes associated with iTreg cells (e.g., Il10rb, Tgfbr3) displayed reduced accessibility in switched Foxp3^+^ Treg-like cells compared to iTreg cells (**Fig. 5c**), while the Th17-related gene (e.g., Rora) was more open, perhaps reflecting relatedness to Th17 cells (**Fig. 5c**). Individual chromatin region traces depict these differences, where switched Foxp3^+^ Treg-like display reduced accessibility for Th17 genes (e.g., Rorc, Il17) but enhanced accessibility for genes associated with iTregs (e.g., Foxp3) (**Fig. 5d**).

**Fig. 5.**
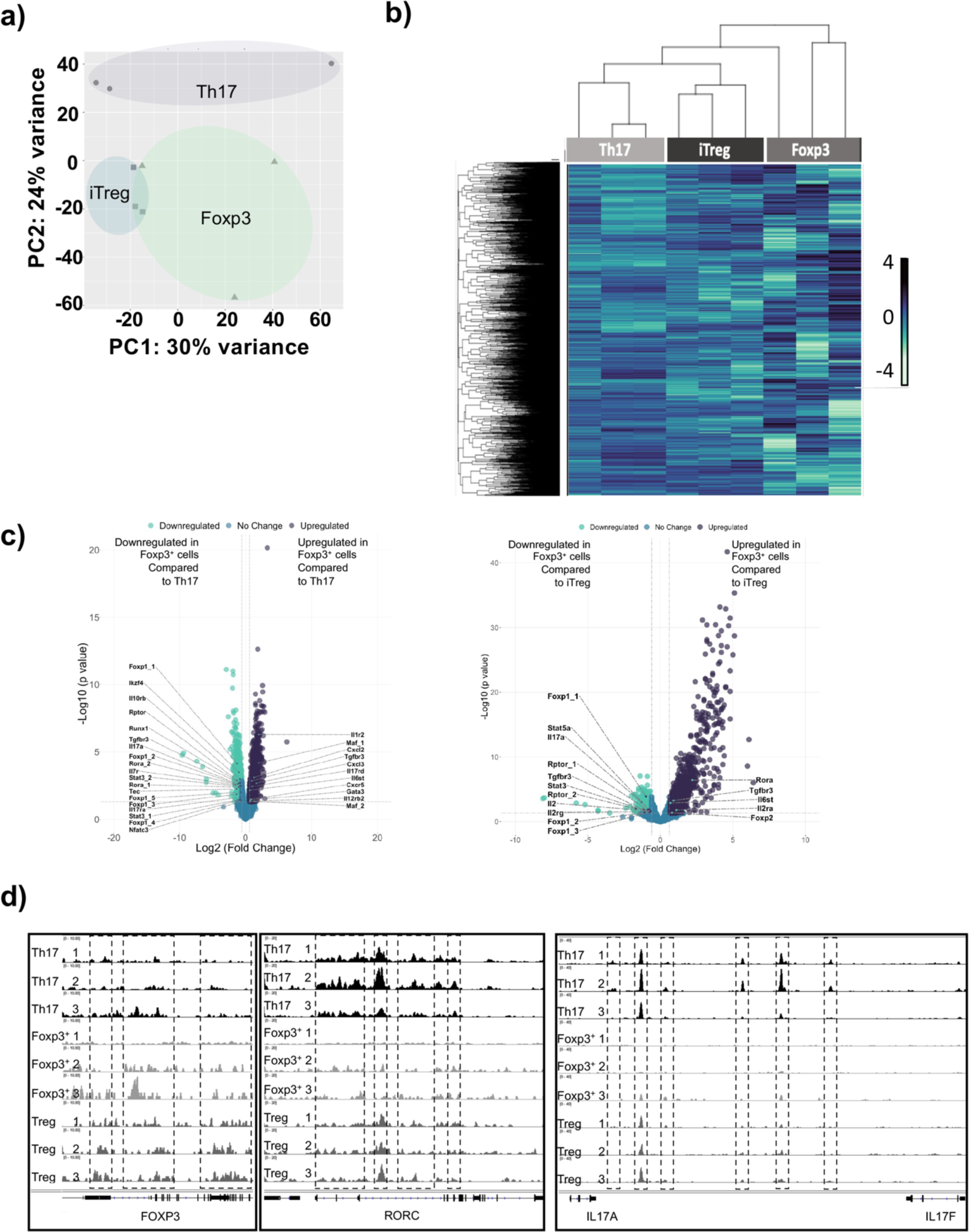
Switched Foxp3^+^ Treg-like cells generated under ITK inhibition have chromatin accessibility profiles distinct from Th17 and iTregs. Switched Foxp3^+^ Treg-like cells were compared with *in vitro* generated Th17 and iTregs by ATAC-Seq analysis. The chromatin accessibility profiles were compared via (**a**) PCA analysis, (**b**) heatmap of global differential peaks in chromatin accessibility, and fold changes of global differential peaks in chromatin accessibility for switched Foxp3 cells compared to (**c**) Th17 cells and iTregs. (**d**) Tracks indicate chromatin areas chromatin accessibility for Foxp3, RORC, IL17A and IL17F in Foxp3^+^ Treg like cells, compared to Th17 cells and iTreg cells.

### Enhanced calcium signaling prevents the ITK dependent switch response to Foxp3^+^ Treg-like cells generated

Differentiation of naïve CD4^+^ T cells into Th17 cells requires both presence of cytokines and TCR activation. An important component downstream of TCR activation is calcium signaling, and we have previously shown that increased intracellular calcium is able to rescue the development of Th17 cells in the absence of ITK (*14*). To determine whether this pathway downstream of ITK is important for the ITK activity dependent switch response, we used ionomycin to enhance cytosolic calcium levels during Th17 differentiation conditions in the presence of ITK inhibition. In the absence of ITK expression (using naïve *Itk^-/-^* CD4^+^ T cells), Th17 differentiation is prevented, and this is rescued by ionomycin treatment, confirming the role of the calcium pathway in Th17 differentiation downstream of ITK (**Fig. 6a**). In addition, there is a switch to Foxp3^+^ Treg-like cell generation, and notably, this switch is prevented by increasing intracellular calcium, indicating a critical role for the calcium pathway in tuning this switch (**Fig. 6a**). To examine if this is also observed with ITK inhibition, we enhanced cytosolic calcium levels with ionomycin during Th17 differentiation conditions in the presence of ITK inhibitor (CPI-818). Increasing intracellular calcium enhanced Th17 differentiation in the absence of ITK inhibition, and importantly, rescued Th17 differentiation in the presence of ITK inhibition (**Fig. 6b**). Together these results indicate that increases in intracellular calcium signaling promotes generation of Th17 cells under the Th17 differentiation condition, and suppresses the generation of switched Foxp3^+^ Treg-like cells in the absence of ITK. Furthermore, while inhibiting ITK’s activity led to increased differentiation of iTregs as we have previously reported (*17*), when ITK is absent, or its ITK’s activity is inhibited (**Fig. 6c,d**). We also noted that inducing increases in intracellular calcium with ionomycin led to suppression of induced Foxp3^+^ Treg cells differentiation, with some observed differentiation of Th17 cells, when naïve CD4^+^ T cells were cultured under iTreg conditions, suggesting that the negative regulation of Treg differentiation, and more broadly, tuning of Th17/Treg differentiation by ITK also travels partly via calcium signaling.

**Fig. 6.**
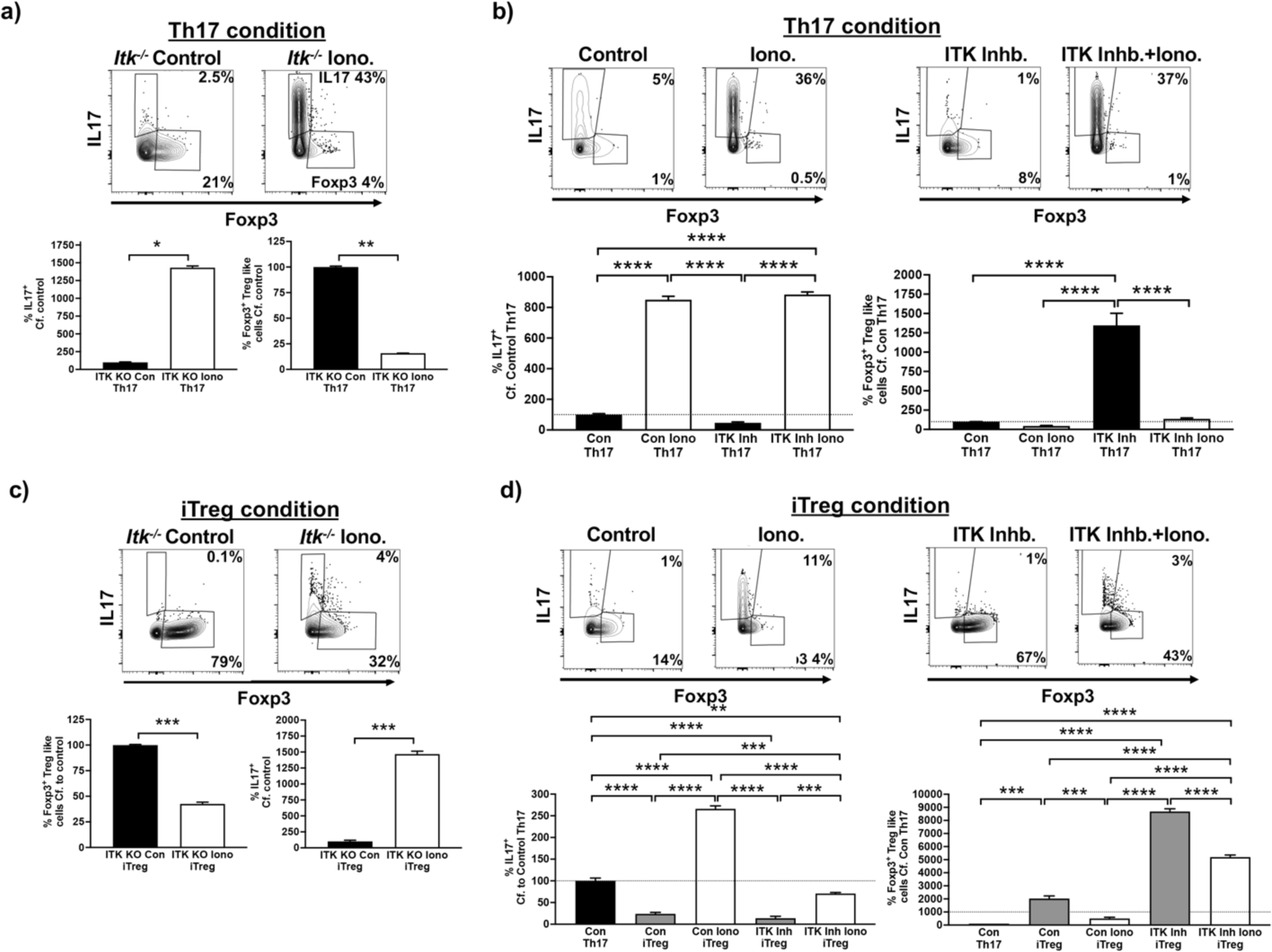
Enhancing calcium signaling prevents the switch to Foxp3^+^ Treg-like cells during ITK inhibition. (**a**) Naïve *Itk^-/-^* IL17A-GFP/Foxp3-RFP CD4^+^ T cells were activated under Th17 differentiation conditions in presence of ionomycin or DMSO control, followed by analysis of percentage of GFP^+^/IL17^+^ cells and percentage of RFP^+^/Foxp3^+^ Treg-like cells. Representative flow plots (top), Quantified (bottom). (**b**) Naïve WT IL17A-GFP/Foxp3-RFP CD4^+^ T cells were activated under Th17 differentiation conditions in presence of ionomycin or DMSO control, with or without ITK inhibitor CPI-818, followed by analysis of percentage of GFP^+^/IL17^+^ cells and percentage of RFP^+^/Foxp3^+^ Treg-like cells. Representative flow plots (top), Quantified (bottom). (**c**) Naïve *Itk^-/-^*IL17A-GFP/Foxp3-RFP CD4^+^ T cells were activated under iTreg differentiation conditions in presence of ionomycin or DMSO control, followed by analysis of percentage of GFP^+^/IL17^+^ cells and percentage of RFP^+^/Foxp3^+^ Treg-like cells. Representative flow plots (top), Quantified (bottom). (**d**) Naïve WT IL17A-GFP/Foxp3-RFP CD4^+^ T cells were activated under iTreg differentiation conditions in presence of ionomycin or DMSO control, with or without ITK inhibitor CPI-818, followed by analysis of percentage of GFP^+^/IL17^+^ cells and percentage of RFP^+^/Foxp3^+^ Treg-like cells. Representative flow plots (top), Quantified (bottom). Mean ± SEM, T test was performed for statistical significance where * p ≤ 0.05, ** p ≤ 0.005, *** p ≤ 0.001 and **** p ≤ 0.0001, 3 independent experiments.

### ITK dependent switch to Foxp3^+^ Treg-like cells involve deficits in molecules involved in mitochondrial oxidative phosphorylation, and reduced expression of BATF

Previous studies have shown that a select set of genes are associated with pathogenic Th17 cells and non-pathogenic Th17 cells (*25, 26*). By comparing the expression levels of these select genes, we find that the level of expression by switched Foxp3^+^ Treg-like cells are distinct from Th17 cells and instead resemble iTregs (**Fig. 7a**). Several metabolic changes are also known to be associated with Th17 cell development, and thus we compared the expression level of select genes involved in these pathways during Th17 cell development (*25, 26*). For genes that are associated with tricarboxylic acid (TCA) cycle activity and mitochondrial function, switched Foxp3^+^ Treg-like cells display expression levels distinct from Th17 cells and instead resemble iTregs (**Fig. 7a**). In contrast for genes that are associated with the hypoxia-induced factor (HIF) pathway and glycolysis, switched Foxp3^+^ Treg-like cells resemble Th17 cells but not iTregs in the level of expression of these markers (**Fig. 7a**). Therefore the switched Foxp3^+^ Treg-like cells seem to have an intermediate metabolic phenotype, resembling in part both the Th17 and iTreg metabolic phenotypes. Next we examined the cells for evidence of activation of select molecules involved in the pathways of mitochondrial oxidative phosphorylation (*25*), by estimating the phosphorylation status of S6K and mTOR, shown to be involved in Th17 states. Using flow cytometric analysis of naïve CD4^+^ T cells cultured under Th17 differentiation condition either with or without ITK inhibitor (CPI-818), we find reduced phosphorylated S6K and mTOR expressing CD4^+^ T cells when ITK’s activity was inhibited compared to control conditions, suggesting reduced mitochondrial oxidative phosphorylation upon ITK inhibition (**Fig. 7b**).

**Fig. 7.**
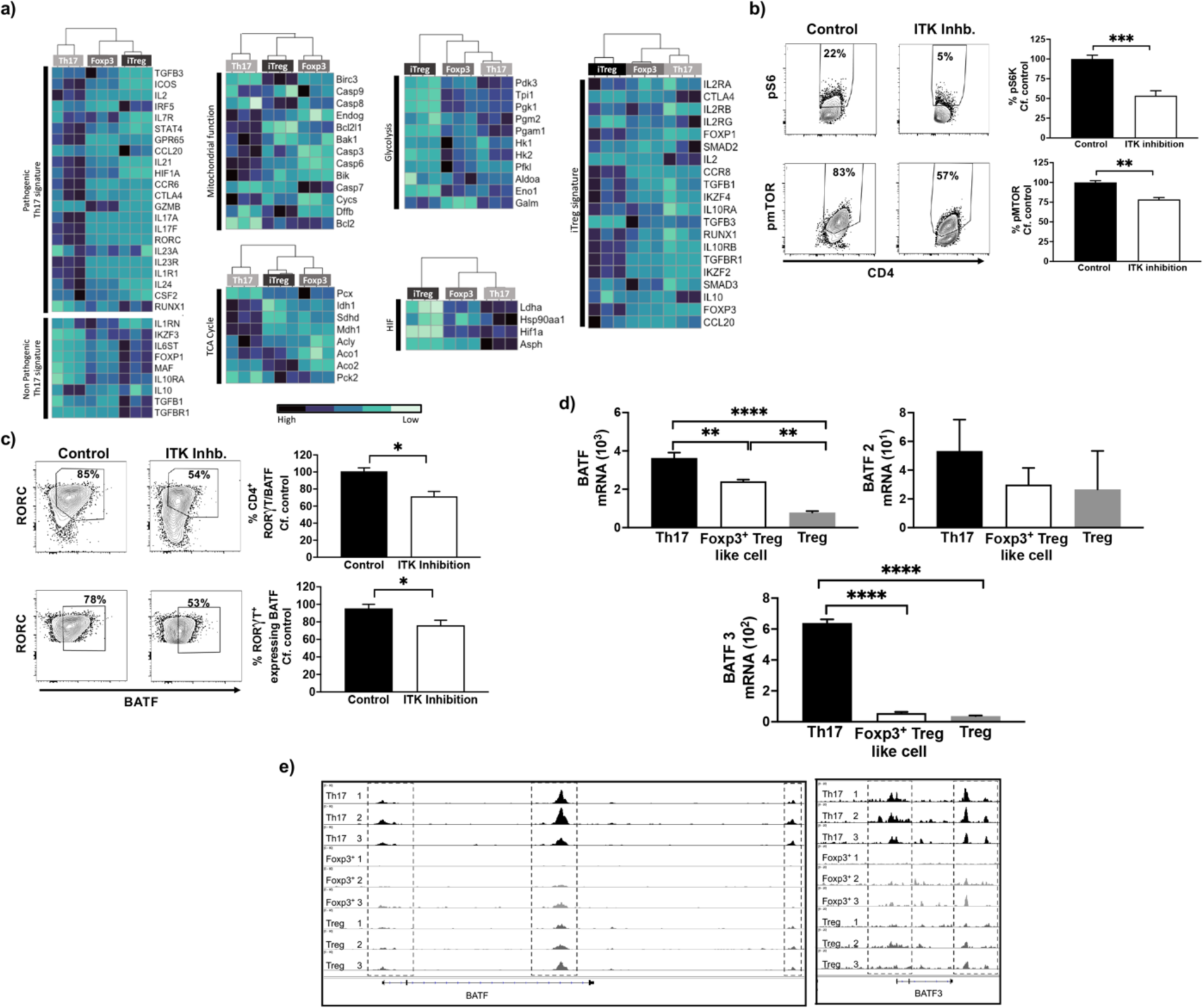
Generation of switched Foxp3^+^ Treg like cells involve altered metabolic pathways and BATF expression. (**a**) The transcriptomes of switched Foxp3^+^ Treg-like cells generated when naïve CD4^+^ T cells are activated under Th17 differentiation conditions in presence or absence of WT ITK inhibitor CPI-818, were compared with signature genes that are associated with pathogenic/non-pathogenic Th17 cells, and genes involved in the TCA cycle, mitochondrial function, HIF1α and glycolysis. (**b**) Flow cytometric analysis of phosphorylation of Ribosomal S6 and mTOR in naïve CD4^+^ T cells are activated under Th17 differentiation conditions in presence or absence of WT ITK inhibitor CPI-818. (**c**) Expression analysis by flow cytometry of BATF in RORC expressing naïve CD4^+^ T cells are activated under Th17 differentiation conditions in presence or absence of WT ITK inhibitor CPI-818. (**d**) Analysis of mRNA expression and (**e**) chromatin accessibility for BATF.

The transcription factor BATF has been identified as a pioneer transcription factor for Th17 differentiation involved in early events during Th17 differentiation (*25*). Our results further indicated reduced overall BATF-expressing CD4^+^ T cells with ITK inhibition, and that RORC expressing cells display reduced BATF expression under Th17 conditions when ITK’s activity was inhibited compared to control (**Fig. 7c**). Similarly we observed reduced mRNA expression and reduced chromatin accessibility for BATF in the switched Foxp3^+^ Treg-like cells compared to the Th17 and iTreg cells (**Fig. 7d,e**). Taken together we demonstrate that ITK inhibition under Th17 differentiation conditions leads to deficits in molecules which mediate mitochondrial oxidative phosphorylation and failure in expression of BATF, which may be involved in the ITK dependent switch to Foxp3^+^ Treg-like cells under Th17 differentiation conditions.

## DISCUSSION

The ability of TCR signals to regulate the differentiation of Th17 and Treg cells has been a focus of intense study, as a better understanding of this process may provide approaches for differential regulation of these T effector subsets in specific immune responses and autoimmune disease. We demonstrate here that the tyrosine kinase ITK tunes a switch between Th17 cells and Foxp3^+^ Treg cells, and that is mediated by calcium signaling, altered cellular metabolism, involving at least in part reduced mTORC signaling with BATF functioning as one of the downstream components. Our findings provide significant mechanistic understanding of how ITK regulates Th17 cells and Treg cells (*14, 19, 27*), and suggest that ITK signaling may control inflammation and anti-inflammation during activation of naïve CD4^+^ T cells. Our work further shows that this ITK mediate tuning of Th17 and Treg cells requires specific signals from ITK during the initial 24-48 hr. period of activation under Th17 conditions, indicating early molecular events regulate the switch response.

In this study, along with expression of Foxp3, we found that switched Foxp3^+^ Treg like cells, resemble iTregs both in terms of phenotypic profile based on the expression of phenotypic Treg markers CD25, CTLA4 and PD1, as well as functionally suppressive function *in vitro*. Foxp3 expression is critical for the development and function of Treg cells, however more recently studies have shown expression of Foxp3 alone is not sufficient for Treg mediated suppressive function (*28, 29*). Recent studies have reported that based on the level of Foxp3 and CD45R expression within human lymphocytes, a heterogenous Treg population could be established where each Treg subset varied in both phenotype and function (*28, 29*). In addition to select phenotypic markers, several studies have utilized comparison of the whole transcriptome and epigenetic landscape to compare T cell subsets. Whereas the transcriptome allowed clear demarcations of phenotypic profile of genes expressed in each T cell subset, investigation into chromatin accessibility status provided a more nuanced view of these cellular states between the different T cell subsets. Comparing the chromatin profile of Foxp3 loci by ChIP Seq for example has revealed that selective activating histone modifications exist in iTregs and not Th1 or Th2 cells, but surprisingly these activating marks were also present in Th17 cells, indicating some similarities in overall chromatin structure of iTregs and Th17 cells (*30*). This previous study also suggested that the chromatin landscape of *in vitro* derived T cells display considerable plasticity, because when studying the loci of the Th2 transcription factor Gata3, it was reported that in order to achieve full activation status and associated histone modifications, two rounds of Th2 polarization were required instead of a single Th2 polarization period (*30*). These observations may explain the discrepancy we observed between transcriptomes and the chromatin accessibility profile. Thus while the switched Foxp3^+^ Treg like cells appear to have attained the transcriptomic profile of iTregs, but further rounds of polarization may be required to induce full transition to iTreg chromatin accessibility profile.

We and others have reported that ITK functions to fine tune the TCR signal strength, being a positive regulator for Th1, Th17 and Tr1 cells but a negative regulator for Th2 and iTregs. Several studies reported that increasing the TCR signal strength enhances differentiation of naïve CD4^+^ T cells into Th17 cells under Th17 polarization conditions but inhibit differentiation of naïve CD4^+^ T cells into iTreg cells under iTreg polarization conditions (*14, 15, 31-34*). Increased TCR signaling is associated with enhanced calcium signaling (*35, 36*) and previously we and others have demonstrated that increasing calcium dependent calmodulin and NFATc1 signaling restores the Th17 differentiation of naïve *Itk^-/-^* CD4^+^ T cells polarized under Th17 conditions (*14, 15, 34*). The findings of the present study showed that enhancing calcium increases using ionomycin enhanced Th17 differentiation in naïve CD4^+^ T cells, but also rescued Th17 differentiation when ITK’s activity was inhibited. Remarkably, this prevented the switch to Foxp3^+^ Treg like cells when ITK’s activity was inhibited under Th17 polarization condition. Furthermore increased calcium not only prevented the switch to Foxp3^+^ Treg like cells with ITK inhibition under Th17 polarization condition (IL6 and TGFβ), but also prevented the increased iTreg differentiation with ITK inhibition under iTreg polarization condition (IL2 and TGFβ), suggesting that the calcium regulated pathway is a key regulator of these two fates.

Calcium signaling was previously reported to control mitochondrial metabolism, such that deletion of stromal interaction molecule 1 (STIM1) in Th17 cells lead to reduced expression of genes for molecules involved in mitochondrial oxidative phosphorylation (*26*). Others have reported the role of enhanced mitochondrial oxidative phosphorylation and glycolysis in Th17 cell differentiation (*11, 12*), which is consistent with our results of increased expression of markers involved in mitochondrial function, oxidative phosphorylation and glycolysis in the Th17 cells. Interestingly we find that the switched Foxp3^+^ Treg like cells generated under ITK inhibition display reduced expression of these markers for mitochondrial function, further emphasizing their altered fate from Th17 cells.

Altered cellular metabolism has been shown to alter T cell differentiation fate via the regulation of the mTOR pathway (*37, 38*), where the inhibition of mTOR in naïve CD4^+^ T cells prevented the generation of IL17A producing Th17 cells and instead lead to generation of Foxp3^+^ iTreg cells, under Th17 polarizing conditions (*37*). In addition several studies have reported reduced mTOR signaling induces iTreg differentiation (*14, 32, 33*), whereas increased mTOR signaling induce Th17 differentiation (*39, 40*). Within Th17 cells, the observed increase in mitochondrial function and oxidative phosphorylation via enhanced mTOR signaling, was further shown to induce expression of the Th17 pioneer transcription factor BATF, allowing maintenance of differentiated Th17 cells (*25*). In contrast the deletion of BATF in naïve CD4^+^ T cells, was reported to increase expression of Foxp3 even under Th17 polarizing conditions (*25, 41*). Our results are therefore consistent with these reports, and further demonstrate that with inhibition of ITK, the switch in T cell fate from Th17 to Foxp3^+^ Treg like cells, under Th17 polarizing conditions, similarly involve the mTOR pathway. These results suggest that with ITK inhibition the reduced mitochondrial function, oxidative phosphorylation and glycolysis, via reduced phosphorylation of mTOR and the mTOR substrate S6K, regulates the switch to Foxp3^+^ Treg like cells. Reduced mTOR signaling via the reduced expression of Th17 transcription factor BATF, serves as a potential mechanism for the ITK dependent switch from Th17 to Foxp3^+^ Treg like cells, under Th17 polarizing conditions.

In conclusion, our results suggest that under Th17 conditions, strong TCR signaling in the presence of ITK activity that leads to increased intracellular calcium, naïve CD4^+^ T cells, results in enhanced mitochondrial function and oxidative phosphorylation, which can activate the mTOR pathway to induce Th17 transcription factor BATF. In the absence of ITK activity, the reduction in expression of molecules involved in mitochondrial function and oxidative phosphorylation, subsequent reduced activity of the mTOR pathways and BATF expression could act as a potential mechanism for the switch in naïve CD4^+^ T cell fate into generating Foxp3^+^ Treg like cells instead. The findings of this study provide greater insight into how ITK controls the Th17/Treg dichotomy, and these findings could have broader implications for immune disorders with an imbalance of Th17/Treg.

## MATERIALS AND METHODS

### Mice

Male and female mice were on the C57BL/6 background aged between 6 to 8 weeks. Mice were housed in the specific pathogen free facilities and all experiments were performed in accordance with the guidelines established by the Office of Research Protection’s Institutional Animal Care and Use Committee at Cornell University. WT, *Itk^-/-^*and ITK*as* IL17-GFP/Foxp3-RFP dual reporter strains were generated by crossing IL17-GFP (B-IL17A-EGFP*^tm1^*, Biocytogen, Worchester, MA) with Foxp3-RFP (C57BL/6-*Foxp3^tm1Flv^*/J, Jackson Laboratory, Bar Harbor, ME) strain (*24*) as previously reported (*16*). CD45.1 congenic (B6.SJL-*Ptprc^a^ Pepc^b^*) and Rag1^-/-^ (B6.129S7-*Rag1^tm1Mom^*) strains were purchased from Jackson Laboratory.

### Flow Cytometry and antibodies

The following antibodies for murine antigens were used labelled as antigen (clone name; catalog number) at a dilution of 1:200 unless indicated otherwise. Viability dye eF506 (65-0866-18) and antibody against CD16/32 (FcBlock) (93; 14016185) were purchased from Thermo Fisher Scientific (Waltham, MA). Antibodies against PD1 (29F.1A12; 135224), mTOR (pSer2448) (MRRBY, 46971842), CD25 (29F.1A12; 135224), CD4 (GK1.5; 100427), CD45.1 (A20; 110716), CD45.2 (104;

109819), CD3ε (145-2C11; 100340) and CD28 (35.51; 102112) were purchased from Biolegend (San Diego, CA). Antibody for S6 (pS235/pS236) (N7-548; 561457) was from BD (San Jose, CA). Flow cytometry data was generated using the Attune Nxt Flow Cytometer (Thermo Fisher Scientific Waltham, MA) and FACS Aria II (BD, San Jose, CA), which was analyzed using FlowJo (Tree Star, Ashland, OR).

### Th17 and iTreg differentiation *in vitro*

Purified naïve CD4^+^ T cells were cocultured with APCs for indicated period of days in RPMI media (10% FBS, 0.5% HEPES, 1 mM Glutamine, 1 mM Sodium Pyruvate, 1 mM non-essential amino acids and 100 U/ml Pen/Strep). Cells were treated with WT ITK inhibitor – CPI-818 (*23*) (Corvus Pharmaceuticals, Burlingame CA), ITK*as* inhibitor – 3MB-PP1 (Cayman Chemicals, Ann Arbor, MI) or DMSO (Sigma) control as indicated. For Th17 differentiation, naïve CD4^+^ T cells were activated by anti-CD3 (2 μg/ml), anti-CD28 (1 μg/ml), in the presence of APCs along with recombinant IL6 (406-ML-025, 10 ng/ml), recombinant TGFβ (240-B-002, 10 ng/ml), and recombinant human IL-2 protein (202-IL-010), all from R&D Systems (Minneapolis, MN) as described (*15*). The iTregs were generated *in vitro* by activating coculture of naïve CD4^+^ T cells and APCs with anti-CD3 (1 μg/ml), anti-CD28 (1 μg/ml), recombinant IL2 (10 μg/ml) and recombinant TGFβ (10 μg/ml). Cells were stained with cell surface markers and/or fixed and permeabilized with the Foxp3/Transcription Factor Staining Kit (eBioscience) with staining for intracellular makers and fixable viability dye eF506 for live/dead cell exclusion to analyze by flow cytometry. Where indicated, cells were also stained with surface markers to sort purify (>95% purity) population of CD4^+^ IL17-GFP^+^ Th17 cells and CD4^+^ Foxp3-RFP^+^ Treg-like cells by BD FACS Aria II.

### *In vitro* suppression assay

Foxp3^+^ Treg-like cells generated during WT ITK inhibition (CPI-818) under Th17 conditions, and iTregs generated during activation under iTreg conditions, were stained with cell surface markers and sort purified (>95% purity) to obtain CD4^+^ Foxp3-RFP^+^ Treg-like cells and CD4^+^ Foxp3-RFP^+^ iTregs. Sort purified naïve responding T cells were labelled with CFSE proliferation dye (Invitrogen) and cocultured with Foxp3^+^ Treg-like cells or iTregs in presence of anti-CD3 (1 μg/ml), followed by analysis using flow cytometry.

### RNA-and ATAC-Sequencing

Total RNA was extracted from sort purified *in vitro* generated Th17, Foxp3^+^ Treg-like cells and iTregs were by TRIzol reagent (Invitrogen). The RNA library was generated using the NEB Ultra II Directional RNA Library Prep Kit and quantified using Qubit Bioanalyzer. RNA sequencing was performed on an Illumina NextSeq500 at 75bp reads and 20 million reads per sample (Transcriptional Regulation and Expression Facility, Cornell University) as previously described (*25*). RNA sequencing data was demultiplexed by BCL2FASTQ2 and FASTQC performed. RNA-Seq data was mapped to the mm10 genome using STAR aligner and raw counts obtained using HTSeq. Counts were normalized and differentially expressed genes identified using DESeq2 (FDR<0.05). Gene set enrichment analysis (GSEA) was performed using software developed by Broad Institute (*26, 27*) and heat map generated using R Studio (Boston, MA) and Heatmapper (*28*). Data was compared to published RNA-Seq data from GEO dataset GSE158703 with the indicated T helper cell subsets.

For ATAC-Seq, Foxp3^+^ Treg-like cells and iTregs generated *in vitro* were sort purified and frozen in 10% DMSO in cell culture media. Nuclei was permeabilized (10 mM Tris-HCl, pH 8.0, 10 mM NaCl, 2 mM Mg Acetate) and lysed (10 mM Tris-HCl, pH 8.0, 10 mM NaCl, 2 mM Mg Acetate, 6 mM CaCl_2_, 0.2% Ipegal, 0.016% Tween20, 600 mM Sucrose) as per instructions from Transcriptional Regulation and Expression Facility, Cornell University. The lysed nuclei were provided to the Transcriptional Regulation and Expression Facility, Cornell University for transposition reaction (*25*) by the Transcriptional Regulation and Expression Facility, Cornell University. DNA library generated was quantified using Qubit BioAnalyzer and sequencing performed on Illumina NextSeq500 at 75bp reads and 20 million reads per sample followed by FASTQC analysis for quality control. ATAQ-Seq data was aligned to mm10 genome using Bowtie, ATAC-Seq peaks were identified by MACS2 and promoter region associated peaks were identified by bedtools as described in (*29*), by the Transcriptional Regulation and Expression Facility, Cornell University. Analysis was performed in R Studio (Boston, MA) and using the UCSC genome browser (University of California Santa Cruz).

### Statistical analysis

Students T test and One way ANOVA was performed for comparison between samples with p<0.05 considered as statistically significant using GraphPad Prism (San Diego, CA).

## Acknowledgments

We thank Amie Redko for animal care, members in the August lab for comments and feedback, Dr. James Janc (Corvus Pharmaceuticals) for CPI-818, and Dr. Jen Grenier of the RNA Sequencing Core for guidance. This work was supported in part by grants from The National Institutes of Health (AI120701 and AI138570 to AA), (AI129422 to AA and WH), and a HHMI Professorship to AA. The National Institutes of Health to The RNA Sequencing Core (U54 HD076210)

## Author contributions

Conceptualization: AA, WH; Methodology: OA, WH; Investigation: OA, WH; Visualization: OA, WH, AA; Funding acquisition: AA, WH; Project administration: AA; Supervision: AA; Writing – original draft: OA, AA; Writing – review & editing: OA, WH, AA.

## Competing interests

AA declares research funding from the 3M Company.

## Data and materials availability

RNA-Sequencing and ATAC-Seq data will be deposited in public databases. All other data are available in the main text or the supplementary materials.

