## Supplemental Data for "ITK signaling regulates a switch between T helper 17 and T regulatory cell lineages via a calcium-mediated pathway"

### Supplementary Materials

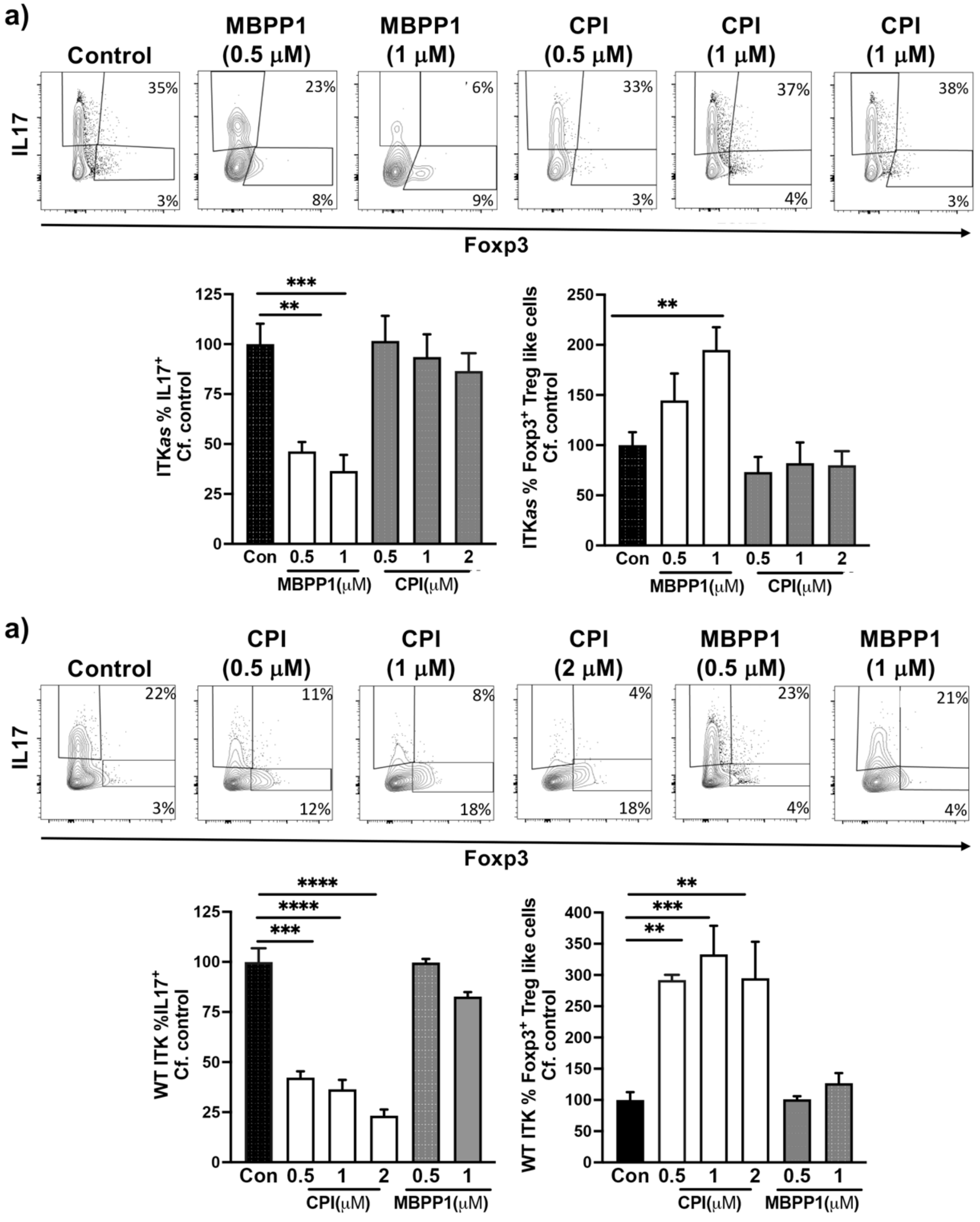


**Fig. S1. The effects of MBPP1 and CPI-818 on Th17 cells is specific.** (**a**) Naïve ITK*as* IL17A-GFP/Foxp3-RFP $CD4^+$ T cells were activated under Th17 differentiation conditions (anti-CD3/28, IL6 and TGFβ) in presence of ITK*as* inhibitor 2MBPP1, CPI-818, or DMSO control as indicated, followed by flow cytometric analysis for percentage of $GFP^+/IL17^+$ cells and $RFP^+/Foxp3^+$ Treg-like cells. (**b**) Naïve WT IL17A-GFP/Foxp3-RFP $CD4^+$ T cells were activated under Th17 differentiation conditions in presence of ITK*as* inhibitor 2MBPP1, CPI-818, or DMSO control as indicated, followed by flow cytometric analysis for percentage of $GFP^+/IL17^+$ cells and

$RFP^+$/$Foxp3^+$ Treg-like cells. Mean ± SEM, Student's T test was performed for statistical significance where * $p \leq 0.05$, ** $p \leq 0.005$, 3 independent experiments.
